# Genetic associations with ratios between protein levels detect new pQTLs and reveal protein-protein interactions

**DOI:** 10.1101/2023.07.19.549734

**Authors:** Karsten Suhre

## Abstract

Protein quantitative trait loci (pQTLs) are an invaluable source of information for drug target development as they provide genetic evidence to support protein function, suggest relationships between *cis*- and *trans*-associated proteins, and link proteins to disease where they collocate with genetic risk loci for clinical endpoints. Using the recently released Olink proteomics data for 1,463 proteins measured in over 54,000 samples of the UK Biobank we identified and replicated 4,248 associations with 2,821 ratios between protein levels (rQTLs) where the strengths of association at known pQTL loci increased by up to several hundred orders of magnitude. We attribute this increase in statistical power (p-gain) to accounting for genetic and non-genetic variance shared by the two proteins in the ratio pair. Protein pairs with a significant p-gain were 7.6-fold enriched in known protein-protein interactions, suggesting that their ratios reflect biological links between the implicated proteins. We then conducted a GWAS on the 2,821 ratios and identified 2,527 novel rQTLs, increasing the number of discovered genetic signals compared to the original protein-only GWAS by 24.7%. At examples we demonstrate that this approach can identify novel loci of clinical relevance, support causal gene identification, and reveal complex networks of interacting proteins. Taken together, our study adds significant value to the genetic insights that can be derived from the UKB proteomics data and motivates the wider use of ratios in large scale GWAS.

## INTRODUCTION

Large-scale studies of the blood circulating proteome leverage the natural variation in the general population to identify genetic and non-genetic factors that control blood protein levels ^1^. Of particular interest for drug development are genome-wide association studies (GWAS) that identify protein quantitative traits (pQTLs), as they provide genetic evidence for a causal effect of the underlying variant – and hence the affected gene(s) – on the levels of the associated protein(s) and their physiological effects. In cases of *cis*-pQTLs, where the genetic variant is located in proximity of the gene coding for the associated pQTL protein, the effect is most likely through a causal variant that modifies transcription, translation or stability of the *cis*-encoded protein. More complex, but also more rewarding in terms of potential biological insights, are *trans*-pQTLs, as they suggest direct or indirect protein-protein interactions between the – presumably causal – *cis-*encoded protein and the associated *trans-*protein, which can extend into larger networks when multiple proteins are associated with a same variant and ideally also clinical endpoints of interest.

Such genetics-driven insights are of highest value to pharmaceutical companies as they can inform drug target discovery and validation, generate hypotheses on modes of action, and suggest biomarkers for target engagement and efficacy. Early successes of pQTL studies ^2–5^ led to the creation of the UKB PPP consortium, a pre-competitive consortium of 13 biopharmaceutical companies that funded the measurement of over 54,000 UKB samples on the Olink Explore 1536 affinity proteomics platform. Olink uses a dual antibody binding technique, termed proximity extension assay (PSA), to quantify the abundance of almost 1,500 blood circulating proteins (**Supplementary Table 1**). The UKB PPP consortium recently published first results from a GWAS that identified over 10,000 pQTLs using this platform ^6^. The Olink proteomics data itself has been released in April 2023 to the public and can be accessed and analyzed using the DNAnexus UKB RAP platform (<u>ukbiobank.dnanexus.com</u>). Our aim in this paper is to explore new methods to enhance pQTL discovery and interpretation, using this exceptional and freely available data set.

We and others previously developed analysis strategies for GWAS with metabolomics data ^7, 8^, a field that is similar in many ways to that of pQTL studies. In particular, we showed that partial correlations between metabolites can reconstruct metabolic networks ^9, 10^ and that the hypothesis-free testing of all ratios between metabolites can substantially strengthen the association signals, in several cases elevating genetic loci out of the background noise ^11, 12^. Both approaches are related in that they identify biological relationships between individual molecules through their shared genetic and non-genetic variance, which can then be integrated into larger metabolic networks, such as the atlas of genetic influences of the human metabolome ^13^ and more recent versions thereof ^14^. Previous GWAS with proteomics suggest that Gaussian graphical models (GGMs) built from partial correlations and ratios between protein levels can reveal biologically relevant protein-protein interactions ^5^, but the approach has never been tested at scale.

Here we hypothesize that a GWAS with ratios between protein levels can identify associations and novel links between protein pairs that have not been identified using current GWAS approaches. However, the computational costs of conducting a full-fledged all-against-all ratio GWAS are prohibitive at this point, estimated to several hundred thousand pounds Sterling on the DNAnexus AWS-based platform for a single run of a full-fledged all-against-all ratio GWAS, not considering costs associated with the handling of the generated data. This challenge will be aggravated in the future by the expected increases in proteome coverage.

We therefore take a more economic approach and test genetic associations with ratios between proteins that are partially correlated and therefore more likely to be related through some biological process. For each pQTL reported by the UKB PPP consortium that implicated one of two partially correlated proteins we test the ratio between the levels of these two proteins for association with the pQTL variant. We then conduct a GWAS on those ratios that increased the strength of association at an already known pQTL locus (see flowchart of this study in **Supplementary Figure 1**). We show in the following that by using this approach we could identify novel pQTLs that were not discovered by the standard GWAS with protein levels conducted by the UKB PPP consortium ^6^, and furthermore, that genetic associations with ratios can uncover biologically relevant links between two or more proteins based on their shared genetic and non-genetic variance. We discuss selected cases of biomedical interest and provide an interpretation of why we believe ratios work.

## RESULTS

### Identification of ratio QTLs at established pQTL loci

We quantify the increase in the strength of an association with ratios by the *p-gain*, which is defined as the smaller of the two p-values for the single protein associations divided by that for the ratio association ^11^. A p-gain of 10 is the equivalent of a nominal p-value for a single test, in other words, a p-gain of 10 is expected to be observed by chance in 5% of the cases when ratios between two random proteins are tested. In the following we require Bonferroni levels of significance for p-values and p-gains throughout and refer to protein ratio associations with significant p-gains as *ratio QTLs* (rQTLs). We split the UKB cohort into a discovery set comprising 43,000 individuals and a replication set of 8,700 individuals based on the data-field *“genetic ethnic grouping”* being equal / not equal to *“Caucasian”* (see UKB documentation on data-field 22006), and further limit the analysis to samples collected at baseline.

A total of 179,923 ratio – variant pairs were tested for association, selected as the overlap of 11,936 Bonferroni significant GGM edges (p < 4.7×10^-8^ or |pcor| > 1.76×10^-3^, **Supplementary Table 2**) and 10,248 Bonferroni significant pQTLs (p < 3.4×10^-11^, **Supplementary Table 3**) from the UKB PPP GWAS ^6^. A total of 10,760 ratio associations (5.98%) had a Bonferroni significant p-gain (> 10* 179,923), and of these 4,248 (41.4%) replicated in the genetically “non-Caucasian” cohort (p-gain > 10*10,760). The 4,248 replicated ratio associations covered 2,821 unique protein pairs between 1,001 of the 1,463 (68.4%) proteins assayed on the Olink platform and 926 of the 5,717 (16.2%) genetic variants reported as pQTL variants by the UKB PPP GWAS (**Supplementary Table 4 & 5**). The likelihood of finding a significant ratio association for a protein pair increased with the strength of their partial correlation from around 5 % for uncorrelated proteins to 9% for |pcor| ∼ 0.2 (**Figure 1**), supporting our choice to prioritize GGM protein pairs.

**Figure 1:**
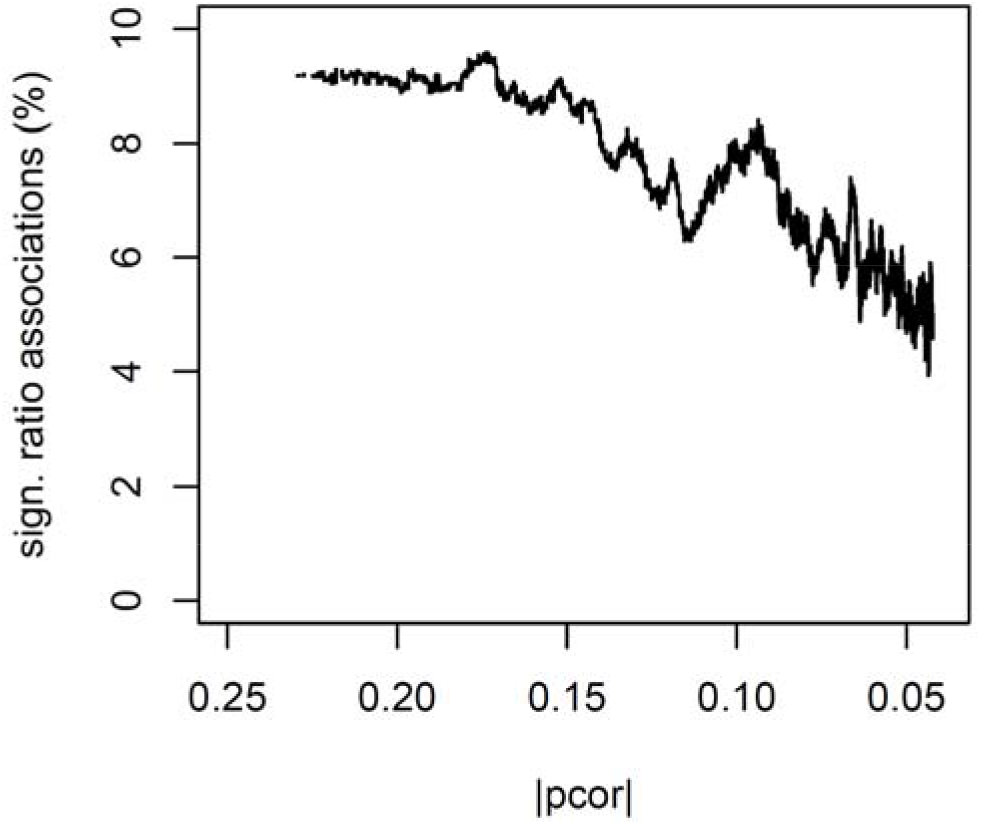
Percentage of Bonferroni significant protein ratio pair associations (p-gain > 10* 179,923) as a function of the partial correlation |pcor| between the protein pair. A moving average with a window size of 10,000 data points was used.

A selection of pharmaceutically relevant rQTLs is provided in **Table 1**, including associations with a ratio between a *cis*- and a *trans*-located protein, where the *cis*-protein is the target of an approved drug (*Tclin* according to Pharos ^15^).

**Table 1:** rQTLs that implicate drug targets in the cis-position. Selected rQTLs that include a ratio between a *cis*- and a *trans*-located protein, with the *cis*-protein being the target of an approved drug (*Tclin* according to Pharos ^15^). The negative log_10_-transformed p-values for the association with the single proteins (-logP.1, -logP.2) and the ratio (-logP.3) and the log_10_- transformed p-gain (logPgain) for the rQTL are reported, the GWAS traits was annotated using PhenoScanner ^16^ (details are in **Supplementary** Table 4).

| Cis Protein | Ratio | Chr | Pos | rsID | -logP.1 | -logP.2 | -logP.3 | logPgain | GWAS traits |
| --- | --- | --- | --- | --- | --- | --- | --- | --- | --- |
| CA6 | CA6/DNER | 1 | 9,034,598 | rs3765963 | 2878.1 | 0.6 | 3235.0 | 356.9 | . |
| LEPR | IL6ST/LEPR | 1 | 66,073,982 | rs10399687 | 0.1 | 225.1 | 270.6 | 45.5 | Blood cell traits |
| SLAMF7 | ICAM3/SLAMF7 | 1 | 160,720,074 | rs11581248 | 3.7 | 2094.1 | 2631.3 | 537.2 | . |
| NECTIN4 | NBL1/NECTIN4 | 1 | 161,049,509 | rs35434391 | 0.4 | 396.6 | 643.6 | 247.0 | Blood cell traits |
| SELP | SELP/VSIR | 1 | 169,563,951 | rs6136 | 349.1 | 0.3 | 663.3 | 314.2 | Activated partial thromboplastin time |
| DPP4 | DPP4/ITGB1 | 2 | 162,930,725 | rs13015258 | 229.0 | 0.9 | 317.5 | 88.5 | . |
| PDCD1 | PDCD1/TNFRSF8 | 2 | 242,801,752 | - | 271.2 | 0.3 | 364.4 | 93.1 | . |
| CD38 | CD38/RELT | 4 | 15,775,851 | rs28703311 | 894.1 | 0.9 | 1030.5 | 136.4 | . |
| PDGFRA | IL6ST/PDGFRA | 4 | 55,139,771 | rs35597368 | 0.6 | 364.8 | 439.3 | 74.4 | Impedance of the legs, Peak expiratory flow |
| F2R | DAG1/F2R | 5 | 76,028,124 | rs168753 | 0.4 | 56.1 | 276.6 | 220.5 | . |
| FLT4 | FLT4/ICAM2 | 5 | 180,057,293 | rs34221241 | 389.3 | 0.6 | 516.6 | 127.4 | . |
| IGF2R | CTSO/IGF2R | 6 | 160,409,894 | rs75474551 | 106.3 | 366.8 | 434.3 | 67.5 | . |
| AKR1B1 | AKR1B1/SUGT1 | 7 | 134,135,621 | rs2229542 | 92.6 | 0.4 | 172.2 | 79.5 | . |
| IMPA1 | IMPA1/TBCC | 8 | 82,583,771 | rs1967328 | 176.4 | 0.9 | 329.5 | 153.0 | . |
| CA1 | CA1/HMBS | 8 | 86,256,210 | rs12544332 | 35.7 | 0.5 | 114.2 | 78.5 | Blood cell traits |
| CA3 | CA1/CA3 | 8 | 86,351,051 | rs2072696 | 6.7 | 109.1 | 387.6 | 278.6 | . |
| CD274 | CD274/EFNA4 | 9 | 5,453,260 | rs822340 | 446.6 | 0.6 | 500.4 | 53.8 | Ulcerative colitis |
| IL2RA | IL2RA/TNFRSF4 | 10 | 6,095,928 | rs12722497 | 1440.8 | 0.1 | 1892.6 | 451.8 | Streptococcal throat infections |
| LAG3 | LAG3/VCAM1 | 12 | 6,885,076 | rs3782735 | 137.5 | 0.6 | 169.1 | 31.6 | . |
| TXNRD1 | NUDT5/TXNRD1 | 12 | 104,707,047 | rs201402862 | 0.4 | 20.3 | 41.8 | 21.5 | . |
| FLT3 | FLT3/FLT3LG | 13 | 28,637,838 | rs9554228 | 34.2 | 36.0 | 53.9 | 17.8 | Age at menarche, Blood cell traits, Body size, mass and fat traits, Rheumatoid arthritis |
| IL4R | CKAP4/IL4R | 16 | 27,327,214 | rs8060025 | 0.3 | 449.7 | 497.6 | 48.0 | Asthma, Eosinophils |
| CA5A | AGXT/CA5A | 16 | 87,927,222 | rs55870502 | 0.9 | 2387.8 | 3738.1 | 1350.3 | Basophils |
| COMT | COMT/HNRNPK | 22 | 19,951,271 | rs4680 | 1008.4 | 0.7 | 3043.4 | 2035.1 | Basal metabolic rate, Systolic blood pressure, Body fat traits |

### Identification of novel rQTLs in a GWAS with ratios

We then conducted a GWAS on the 2,821 ratios using the genotyped UKB data. For each ratio we retained the strongest associations that reached a Bonferroni level of significance of p-value < 5×10^-8^/ 2,821, a p-gain > 10^7^ * 2,821, and that were more distant than one million base pairs from any other significant association with the same ratio. We identified 8,462 ratio-variant pairs with 2,095 unique variants that satisfied this criterion, which corresponds to a discovery per tested ratio of on average three independent GWAS signals with a significant p-gain (**Supplementary Table 6**, **Figure 2, Supplementary Figure 2**). The ratios with the largest number of rQTLs discovered were HBEGF / PDGFA (N=25) and ITGB1BP2 / MITD1 (N=24). A total of 999 proteins were implicated in at least one rQTL, with a median of eight rQTLs per protein. The two most frequently occurring proteins were ITGB1BP2 with 259 rQTLs and EDAR with 237 rQTLs. A total of 2,527 (29.9%) of the 8,462 rQTLs were more distant that 10^6^ base pairs from any pQTL reported by the UKB PPP GWAS for one of the two proteins in the respective ratio and thus represent previously non-reported pQTLs, which corresponds to an increase of 24.7% in genetic signals derived from the UKB PPP Olink data using ratios compared to the standard approach.

**Figure 2:**
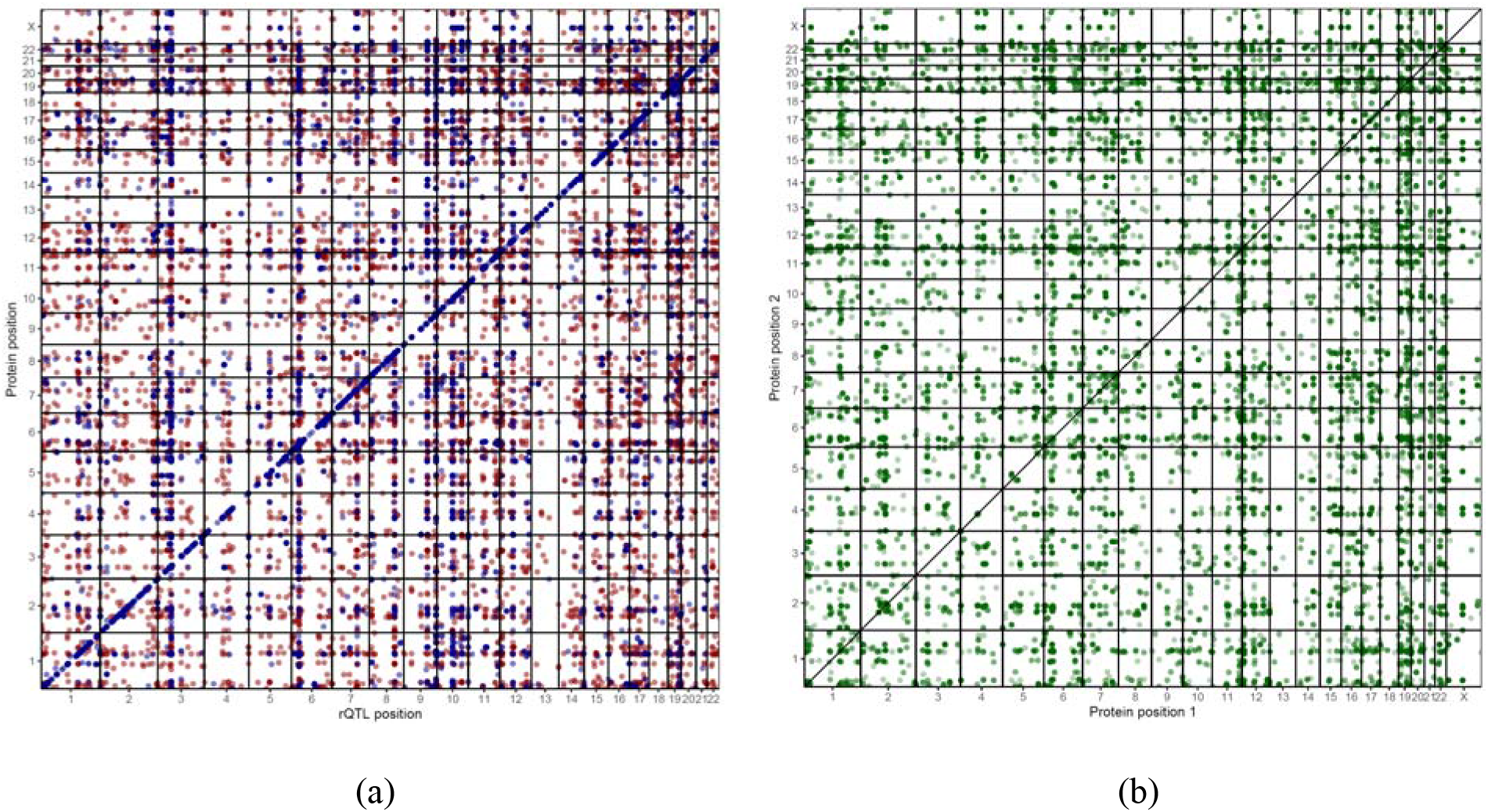
2D-Manhattan plots. (a) The position of the rQTL plotted against the position of the genes coding for the two proteins in the ratio, the stronger of the two single protein associations is in blue, the weaker in red; (b) The positions of the genes coding for the two proteins in an rQTL ratio plotted against each other, darker colors indicate multiple rQTLs with a same ratio.

To investigate whether these rQTLs provided new insights of biomedical interest we annotated the 2,095 rQTL variants identified in this study and the 5,717 pQTL variants reported by the UKB PPP GWAS using PhenoScanner ^16^ for association with 446 distinct GWAS traits (**Supplementary Table 6** and **Supplementary Table 3**, resp.). We identified 322 rQTL variants that were more distant than 10^6^ base pairs from any pQTL variant on the same GWAS trait, implicating 874 rQTLs in a total of 4,700 co-associations with GWAS traits (**Supplementary Table 7**).

These rQTLs provide new evidence to support drug target selection. For instance, rs3764640 associated with the ratio STK11/USP8 (-log_10_(p) = 13.8, log_10_(p-gain) = 10.5). The variant is an intragenic SNP in the *STK11* gene and associated with the presence versus absence of psychosis in Alzheimer’s cases ^17^. STK11 is a serine/threonine-protein kinase and USP8 may play a role in the degradation of activated protein kinases by ubiquitination ^18^, which would explain the significant p-gain for the ratio. This rQTL hence not only supports a role of STK11 in AD pathology, but also provides further insights into the putative underlying biological pathways, suggesting that medicinal modification of STK11 or its phosphorylation targets may affect the AD related phenotypes.

A second example is a region of high LD on chromosome 5 (**Figure 3**) which is a major risk locus for inflammatory bowel disease (IBD) ^19^. The most likely causal gene prioritized by multiple GWAS, based on its function and the presence of an amino acid-changing variant, was *Macrophage Stimulating 1* (*MST1*). However, this view has been challenged, proposing *Glutathione Peroxidase 1* (*GPX1*) as a causal gene instead, supported by biochemical experiments showing that a co-segregating amino-acid variant in *GPX1* reduced the activity of this antioxidant enzyme ^20^. Here we identified 14 ratios between 16 proteins that associated with a significant p-gain at this locus (**Table 2**). Seven were ratios of the *pyruvate kinase, liver and red blood cells* (PKLR) with proteins involved in haemoglobin metabolism, including *hydroxyacylglutathione hydrolase* (HAGH), *hydroxymethylbilane synthase* (HMBS), *Arginase 1* (ARG1), *Biliverdin Reductase B* (BLVRB). The biochemical properties of these genes clearly support a causal role for GPX1 in an oxidative stress related phenotype, likely related to haemoglobin metabolism in red blood cells. However, four of the ratios were with the *cis*-encoded protein *Dystroglycan 1* (DAG1) and several proteins not related to red blood cell metabolism, suggesting the presence of a second, likely independent causal gene at this locus, which would co-segregate with the GPX1 variant due to the high linkage disequilibrium in this region. Whether both pathways are driving factors of the IBD association requires further investigation. Important for our study is that this case exemplifies the kind of insights that can be drawn from using rQTLs and their value for drug target evaluation and hypothesis generation.

**Figure 3:**
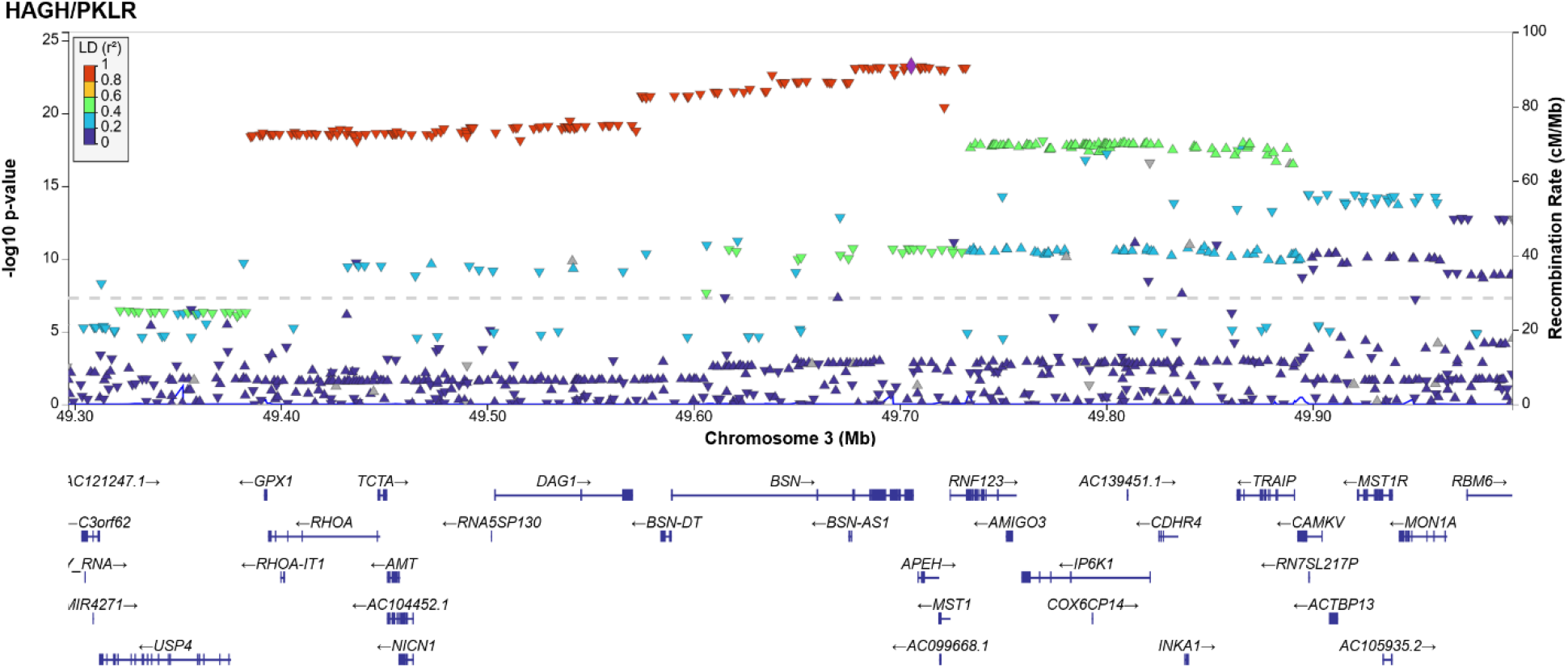
Regional association plot for the association of the HAGH / PKLR ratio at a major IBD locus on Chr3. Plot created using LocusZoom ^43^.

**Table 2:** Novel cis-pQTLs. List of 39 genetic variants that associated with a ratio that involves a protein located less than 1MB from the variant (Cis protein) and that has no cis-pQTL in the UKB PPP GWAS. The negative log_10_-transformed p-values for the association with the single proteins (-logP.1, -logP.2) and the ratio (-logP.3) and the log_10_-transformed p-gain (logPgain) for the rQTL are reported (**details are in Supplementary** Table 6).

| Cis Protein | Ratio | Chr | Pos | rsID | -logP.1 | -logP.2 | -logP.3 | logPgain |
| --- | --- | --- | --- | --- | --- | --- | --- | --- |
| MNDA | MNDA/NCF2 | 1 | 158,788,542 | rs2875712 | 5.7 | 0.3 | 16.9 | 11.2 |
| EPCAM | EPCAM/GPA33 | 2 | 47,773,540 | rs6708696 | 1.6 | 1.3 | 15.5 | 13.9 |
| ANXA4 | ANXA4/LACTB2 | 2 | 70,033,584 | rs2228203 | 11.6 | 3.8 | 232.7 | 221.1 |
| TMSB10 | DBI/TMSB10 | 2 | 85,133,861 | rs13409738 | 0.0 | 8.1 | 36.8 | 28.7 |
| NCK2 | NCK2/PLA2G4A | 2 | 106,452,253 | rs10169998 | 4.6 | 0.0 | 36.4 | 31.8 |
| TGFBR2 | TGFBR2/TNFRSF1A | 3 | 30,729,510 | rs114836705 | 8.2 | 0.4 | 25.0 | 16.9 |
| PPP1R2 | PPP1R2/SNAP23 | 3 | 195,238,559 | rs34950021 | 1.9 | 0.8 | 26.6 | 24.7 |
| CXCL3 | CXCL3/CXCL5 | 4 | 74,797,139 | rs352024 | 0.6 | 514.2 | 1229.3 | 715.1 |
| FYB1 | FYB1/PPP1R12A | 5 | 39,338,358 | rs3822462 | 1.8 | 0.7 | 14.1 | 12.3 |
| DAB2 | DAB2/NCK2 | 5 | 39,427,481 | rs75839063 | 2.5 | 0.0 | 20.3 | 17.8 |
| HBEGF | HBEGF/PDGFA | 5 | 139,720,400 | rs2237077 | 7.6 | 0.8 | 31.1 | 23.4 |
| PDLIM7 | PDLIM7/SRC | 5 | 176,922,643 | rs335428 | 7.5 | 1.0 | 28.9 | 21.4 |
| MPIG6B | MPIG6B/PLXNA4 | 6 | 31,346,193 | rs2507982 | 0.3 | 5.1 | 18.5 | 13.5 |
| MAP3K5 | MAP3K5/MAVS | 6 | 136,888,889 | rs56379668 | 4.0 | 1.9 | 31.7 | 27.7 |
| VTA1 | RWDD1/VTA1 | 6 | 142,641,606 | rs12189801 | 0.6 | 4.2 | 21.1 | 16.9 |
| LAT2 | CLIP2/LAT2 | 7 | 73,780,812 | rs512023 | 4.9 | 4.8 | 59.6 | 54.7 |
| CASP2 | CASP2/NUB1 | 7 | 142,986,684 | rs3181165 | 2.8 | 0.5 | 14.3 | 11.4 |
| PLPBP | PLPBP/RWDD1 | 8 | 37,635,649 | rs7463174 | 8.7 | 1.1 | 37.9 | 29.2 |
| EIF4EBP1<sup>+</sup> | DNPH1/EIF4EBP1 | 8 | 37,884,310 | rs28797500 | 0.5 | 4.3 | 18.4 | 14.1 |
| LYN | KIFBP/LYN | 8 | 56,785,133 | rs6985703 | 0.2 | 5.5 | 18.5 | 12.9 |
| INPPL1<sup>+</sup> | BANK1/INPPL1 | 11 | 72,064,041 | rs79658353 | 0.1 | 1.7 | 13.0 | 11.3 |
| PPME1 | ATG4A/PPME1 | 11 | 73,948,875 | rs79153613 | 0.6 | 9.5 | 30.1 | 20.6 |
| ARHGEF12<sup>+</sup> | ARHGEF12/AXIN1 | 11 | 120,278,477 | rs34172482 | 4.6 | 0.1 | 37.7 | 33.1 |
| IRAG2 | CRACR2A/IRAG2 | 12 | 25,243,115 | rs1908946 | 0.3 | 3.5 | 16.4 | 12.8 |
| METAP2 | EIF4B/METAP2 | 12 | 95,830,338 | rs159853 | 0.6 | 8.9 | 27.4 | 18.5 |
| TRIAP1 | TMSB10/TRIAP1 | 12 | 120,902,007 | rs542407 | 0.3 | 3.6 | 14.7 | 11.1 |
| SRP14 | APEX1/SRP14 | 15 | 40,325,829 | rs6492926 | 0.0 | 10.9 | 25.3 | 14.4 |
| SNAP23 | CD69/SNAP23 | 15 | 42,808,309 | rs73404730 | 0.8 | 1.7 | 21.2 | 19.6 |
| PPIB | MANF/PPIB | 15 | 64,780,971 | rs73452261 | 1.0 | 1.9 | 29.2 | 27.3 |
| MESD | MANF/MESD | 15 | 81,279,706 | rs57967327 | 0.3 | 5.4 | 74.5 | 69.1 |
| CORO1A | CORO1A/TBC1D5 | 16 | 30,147,265 | rs7201780 | 4.9 | 0.0 | 18.2 | 13.3 |
| STX4 | DNMBP/STX4 | 16 | 31,004,812 | rs12445568 | 0.3 | 6.7 | 20.5 | 13.9 |
| AHSP | AHSP/CA2 | 16 | 31,463,216 | rs4889659 | 9.9 | 0.0 | 26.3 | 16.4 |
| VPS53 | SCAMP3/VPS53 | 17 | 400,933 | rs9916346 | 0.3 | 2.9 | 14.7 | 11.8 |
| STK11 | STK11/USP8 | 19 | 1,207,238 | rs3764640 | 3.3 | 0.6 | 13.8 | 10.5 |
| CDKN2D | CDKN2D/TACC3 | 19 | 9,868,278 | rs10420364 | 0.5 | 6.7 | 45.5 | 38.7 |
| CDC37 | CDC37/PLA2G4A | 19 | 10,523,086 | rs10854116 | 4.4 | 0.8 | 47.0 | 42.6 |
| TBCB | MITD1/TBCB | 19 | 36,593,915 | rs61741470 | 0.3 | 9.7 | 130.7 | 121.0 |
| CRKL | CRKL/DBNL | 22 | 21,139,239 | rs117858197 | 1.6 | 0.5 | 34.8 | 33.1 |

<sup>†</sup>This protein has no pQTL in the UKB PPP GWAS

### Discovery of novel cis-pQTLs

Observation of *cis-*pQTLs is considered genetic evidence to confirm the target specificity of the respective affinity binding assay. Sun *et al*. found a *cis-*pQTL for 1,163 (79.5%) of the 1,463 assayed proteins. Here we report 39 additional genetic variants that associated with a ratio that involves a protein located in-cis and a second protein located in-*trans* (**Table 3**). These *cis*-pQTLs became presumably discoverable as the *trans*-proteins in the ratios captured some unidentified shared non-genetic variance, accounting for which lead to the significant p-gains. The corresponding 39 proteins include three Olink targets for which no genetic signal had been found in the UKB PPP GWAS at all (ARHGEF12, EIF4EBP1, INPPL1) and thus provide genetic evidence that the respective antibodies bind their designated targets. Even by using only a subset of all possible ratios, we identified 13% of the 300 cis-pQTLs that were not accounted for so far, increasing confidence in the target specificity of the Olink platform for these proteins. More may be identified in an all-against-all ratio approach.

**Table 3:** Association of selected ratios with rs9858542. Ratios have been arranged such that all associations are negative with the copy number of the G allele of variant 3:49701983:G:A.

| RATIO | Protein name 1 | Protein name 2 | -logP.1 | -logP.2 | -logP.3 | logPgain |
| --- | --- | --- | --- | --- | --- | --- |
| HMBS / PKLR | Hydroxymethylbilane Synthase | Pyruvate Kinase L/R | 3.3 | 8.2 | 45.7 | 37.5 |
| ARG1 / PKLR | Arginase 1 |  | 2.1 | 8.2 | 30.3 | 22.1 |
| BLVRB / PKLR | Biliverdin Reductase B |  | 0.2 | 8.2 | 29.4 | 21.2 |
| HAGH / PKLR | Hydroxyacylglutathione Hydrolase |  | 0.4 | 8.2 | 23.0 | 14.8 |
| LHPP / PKLR | Phospholysine phosphohistidine inorganic pyrophosphate phosphatase |  | 1.2 | 8.2 | 22.0 | 13.8 |
| DNPH1 / PKLR | 2'-Deoxynucleoside 5'-Phosphate N-Hydrolase 1 |  | 0.9 | 8.2 | 20.0 | 11.8 |
| PSMD9 / PKLR | Proteasome 26S Subunit, Non-ATPase 9 |  | 1.4 | 8.2 | 31.6 | 23.4 |
| AHSP / BLVRB | Alpha Hemoglobin Stabilizing Protein | Biliverdin Reductase B | 13.2 | 0.2 | 31.4 | 18.3 |
| AHSP / CA2 | Alpha Hemoglobin Stabilizing Protein | Carbonic Anhydrase 2 | 13.2 | 0.8 | 24.0 | 10.8 |
| HMBS / UBAC1 | Hydroxymethylbilane Synthase | Ubiquitin-associated domain-containing protein 1 | 3.3 | 2.5 | 16.7 | 13.4 |
| CD84 / DAG1 | SLAM family member 5 | Dystroglycan 1 | 0.0 | 15.1 | 34.7 | 19.6 |
| F2R / DAG1 | Coagulation Factor II Thrombin Receptor |  | 0.5 | 15.1 | 62.1 | 47.0 |
| HBEGF / DAG1 | Heparin Binding EGF Like Growth Factor |  | 2.6 | 15.1 | 27.9 | 12.8 |
| SCARF1 / DAG1 | Scavenger Receptor Class F Member 1 |  | 0.1 | 15.1 | 41.9 | 26.8 |

### Refinement of the rQTL loci

For economic reasons, we conducted the GWAS using the genotyped variants only and may therefore have missed variants of interest. For each of the 8,462 rQTLs we therefore refined the associations within +/-500,000 base pairs of the respective lead variant by using the imputed UKB genotype data, both, in the discovery and in the replication cohort. We provide the summary statistics for all 8,462 refined regions on FigShare (doi:10.6084/m9.figshare.23695398). This data can also be used to further refine loci of interest, for instance to identify potentially multiple independent signals using SuSiE ^21^ or to test for colocalization with other traits of interest using coloc ^22^. To visualize individual rQTLs we generated regional association plots for all rQTLs, both in the discovery and the replication cohort (**Supplementary Figure 3**).

We then used coloc ^22^ to ask whether the two proteins in a ratio shared a same genetic signal (Q.12), whether any of the two proteins shared a signal with the ratio (Q.13 and Q.23), and whether the signal for the ratio was shared between discovery and replication cohort (Q.33.repli). **Supplementary Table 6** provides the most likely hypothesis for each of these four questions, together with its posterior probability. In 7,414 (87.6%) of the 8,462 cases at least one of the proteins shared a genetic signal with the ratio (Q.13 = H4 or Q.23 = H4), in 1,305 (15.4%) cases both proteins shared a signal with the ratio (Q.13 = H4 and Q.23 = H4), and in 489 (5.8%) cases there was no signal detectable for either of the two proteins alone (Q.13 = H2 and Q.23 = H2 and Q.12 = H0). A total of 6,775 of the 8,462 rQTLs (80.1%) shared a genetic signal between discovery and replication cohort (Q.33.repli = H4).

For each rQTL region we designated the variant with the strongest association with the ratio in the discovery cohort as the lead variant and asked whether the association on this variant replicated. Requiring in the discovery cohort p-value < 5×10^-8^/2821 and p-gain > 10*10^6^*2821 and in the replication cohort p-value < 0.05/8462 and p-gain > 10*8462, we identified 4,181 rQTLs (49.4%) that satisfied this stringent Bonferroni significance criterion. Considering that 80.1% of the rQTLs shared a same genetic signal between discovery and replication cohort, it is likely that more rQTLs can be replicated when more samples become available.

### Why do ratios work and what do they represent?

With P_1_ and P_2_ representing the levels of two blood circulating proteins (we suppress the indices of the individual samples) we can fit two linear models to the log-scaled protein levels by selecting parameters α*_i_*, β*_i_*, and γ*_i_* such that they minimize the square of the non-explained variance ε*_i_* in the following equation:

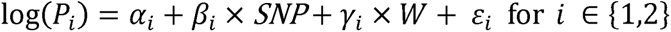

*SNP* represents the number of effect alleles (0, 1, 2) of a given genetic variant in a given sample and *W* denotes some non-identified non-genetic variance that is shared by both proteins. Using the identity log(A/B) = log(A) – log(B), the ratio can then be written as:

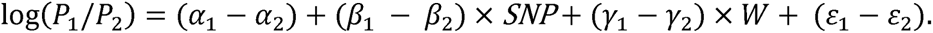

As the significance level (p-value) of the association with the variant depends on the proportion of the variance that is explained by the genetic term (*SNP*) compared to the remaining variance (W+ E), the strength of the association with the ratio can increase under two conditions: (1) when β*_1_* and β*_2_* have opposite signs, or (2) when γ*_1_* and γ*_2_* are of comparable size and non-zero.

β*_1_* and β*_2_* having opposite signs implies that the genetic variant increases the levels of one protein while decreasing those of the other (**Figure 4**). When working with metabolites, this situation can occur when the ratio represents a substrate-product pair of an enzyme who’s efficacy is affected by the genetic variant. Many such cases have been reported ^12, 13, 23^. For proteins, a possible scenario is a genetic variant that increases the expression of one protein which acts as a suppressor of a second protein. One example is the association of rs1065853, which is in LD with coding SNP rs7412 in *APOE*, and the ratio between LDLR and PCSK9 protein levels (log_10_(p-gain) = 67.2). PCSK9 binds LDLR and targets it for degradation ^24^. PCSK9’s availability to degrade LDLR in turn is limited by binding to apolipoprotein B ^25^, the levels of which are associated with rs7412 in *APOE*.

**Figure 4:**
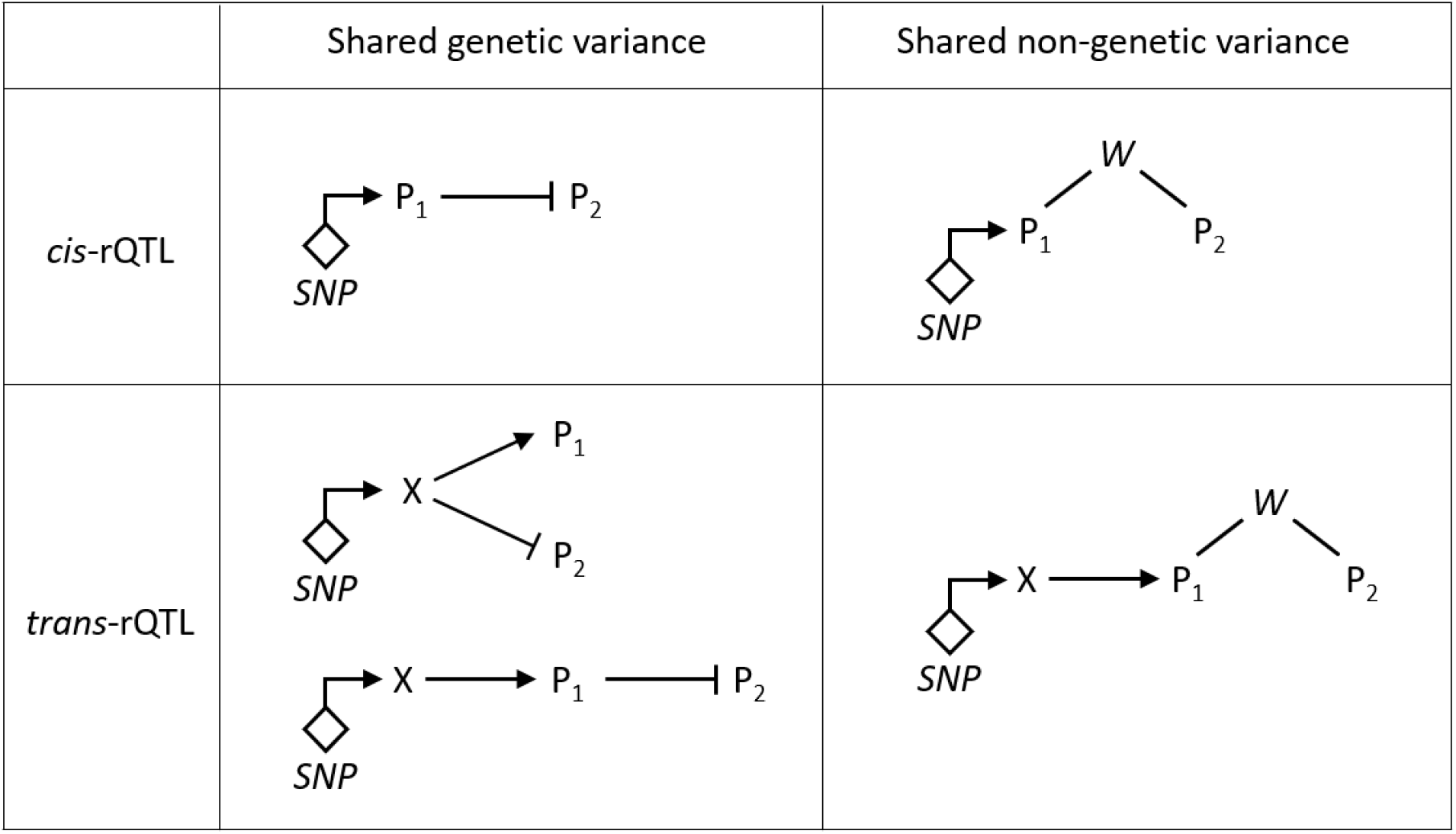
Possible scenarios that can lead to a significant p-gain in a ratio association. P1 and P2 are the proteins in the ratio that associates with the genetic variant *SNP*, X is the causal cis-encoded protein in the case of a trans-rQTLs; *W* denotes some unidentified shared non-genetic variance.

If only one of the proteins is affected by the genetic variant, then observing a significant p-gain implies that γ*_1_* and γ*_2_* must be of comparable size and non-zero, and the association with the ratio indicates the presence of some non-genetic variance that is shared by both proteins. For instance, ITGB1BP2 has only five trans-pQTLs in the UKB PPP GWAS of moderate effect size, but occurs with 26 different ratios in 259 rQTLs in our GWAS, the strongest with a log10(p-gain) = 1115.5 for the association of rs4680 with the ratio COMT / ITGB1BP2 at the *COMT* gene locus. The association of ITGB1BP2 with rs4680 was not significant (p = 0.69). The ratio with the largest number of rQTLs in the GWAS was ITGB1BP2 / MITD1, which had 24 rQTLs compared to only 2 pQTLs for MITD1 in the UKB PPP GWAS (**Figure 5**). Intriguingly, both proteins are highly correlated (Pearson r^2^ = 0.86), suggesting that their correlation is driven by some shared, yet not identified factor. The strongest correlations with one of the clinical biochemistry and blood traits available in UKB was with platelet count (r^2^ = 0.12 with ITGB1BP2, r^2^ = 0.10 with MITD1, and r^2^ = 0.025 with the ratio), which are too weak to explain the full correlation between both proteins, suggesting that a driving factor for this association is related to some more specific, probably blood cell type related trait that is not readily available in the UKB phenotype dataset.

**Figure 5:**
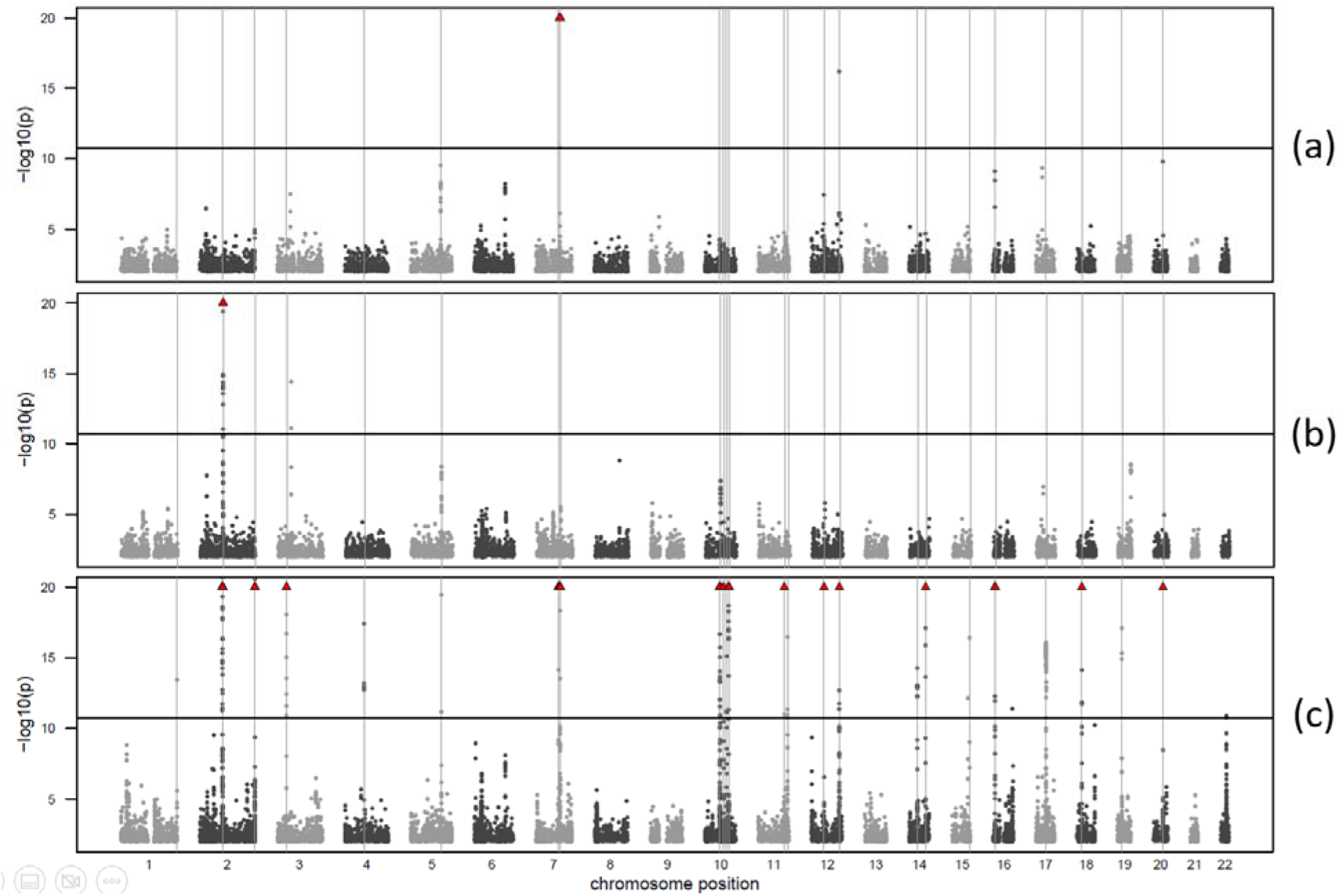
Example of a ratio that leads to the discovery of novel signals. Manhattan plots for the GWAS with (a) ITGB1BP2, (b) MITD1, and (c) the ratio ITGB1BP2 / MITD1. Associations with p-values exceeding 10^-20^ are indicated by red triangles. Vertical lines indicate 24 Bonferroni significant (p < 5×10^-8^/2821) rQTLs for the ratio. Manhattan plots for all 2,821 ratio GWAS are available as online as explained in **Supplementary** Figure 1.

There are thus multiple possible causes that can lead to a significant p-gain in a ratio association, as schematized in **Figure 4**, some rQTLs revealing the presence of shared genetic variance while others suggest the proteins in the ratio being linked through some shared non-genetic processes.

### What can be learned from rQTLs?

To evaluate the enrichment in protein pairs that were linked through significant ratio associations and/or GGM edges we used the STRING database of protein-protein interactions (**Supplementary Table 5**). Of the 2,281 protein pairs, 168 pairs (6.0%) had a protein-protein interaction link in the STRING database with a high confidence score (> 0.7), while random pairs between these proteins had on average only 22.1 links (s.d. = 4.2, based on 100 samplings), which corresponds to a 7.6-fold enrichment. For comparison, of the 11,936 protein pairs linked through significant GGM edges, 465 pairs (3.9%) had a protein-protein interaction reported in TRING, while random sampling yielded an average of 89.4 (s.d. = 9.4), which corresponds to a 5.2-fold enrichment for proteins linked through GGM edges alone.

Using cytokine and cytokine-receptor annotations from CytokineLink ^26^ we identified 39 protein pairs with an rQTL that involved cytokine-cytokine pairs, 15 receptor-receptor pairs, and 9 cytokine-receptor pairs, seven of which were known and two were new (CSF1:LTBR and CXCL9:TNFRSF9). CytokineLink predicted 1,542 cytokine-cytokine interactions between 77 cytokines from the Olink platform (out of 77*76/2 = 2,926 possible interactions). The number of cytokine pairs that were involved in rQTLs was significantly enriched (30 out of 39 compared to 1,512 out of 2,887 with no rQTL, p < 0.002, Fisher exact test).

When multiple proteins associate in rQTLs with a same variant, networks of related proteins can be constructed. Here we present an example of how a single pQTL around a pharmaceutically interesting protein can be extended into a network of potentially interacting proteins. NFATC1 (Nuclear Factor Of Activated T Cells 1) is key transcription factor and regulator of the immune response ^27^ and a molecular target for immunosuppressive drugs such as cyclosporin A ^28^. NFATC1 has been implicated in the pathogenesis and targeted therapy of hematological malignancies ^29, 30^. rs657693 is a *cis-*pQTL for NFATC1 in the UKB PPP GWAS and one of only two genetic association for this protein. Here we identified rs657693 as an rQTL for the ratio of NFATC1 with 16 other proteins (AXIN1, BACH1, BANK1, BCR, CASP2, CD69, EIF4G1, FADD, FOXO1, IKBKG, INPPL1, IRAK1, LBR, PTPN6, SPRY2, TJAP1, **Table S3**), none of which had a significant association with this variant alone. Nine of the 16 proteins had a second replicated rQTL in a ratio with NFATC1 elsewhere in the genome, and in all of these cases the other protein in the ratio was the driving pQTL, with five of them being cis-pQTLs (AXIN1, BANK1, FOXO1, SPRY2, TJAP1). Our GWAS identified additional rQTLs, including cis-rQTLs for BCR (rs713617), CD69 (rs7309767), and FADD (rs7939734). Using IPA we identified a number of functional links between these proteins, generating a network of proteins that can now be linked through genetic evidence and rQTLs to the NFATC1 locus, potentially supporting the development of new immunosuppressive drugs (**Figure 6**).

**Figure 6:**
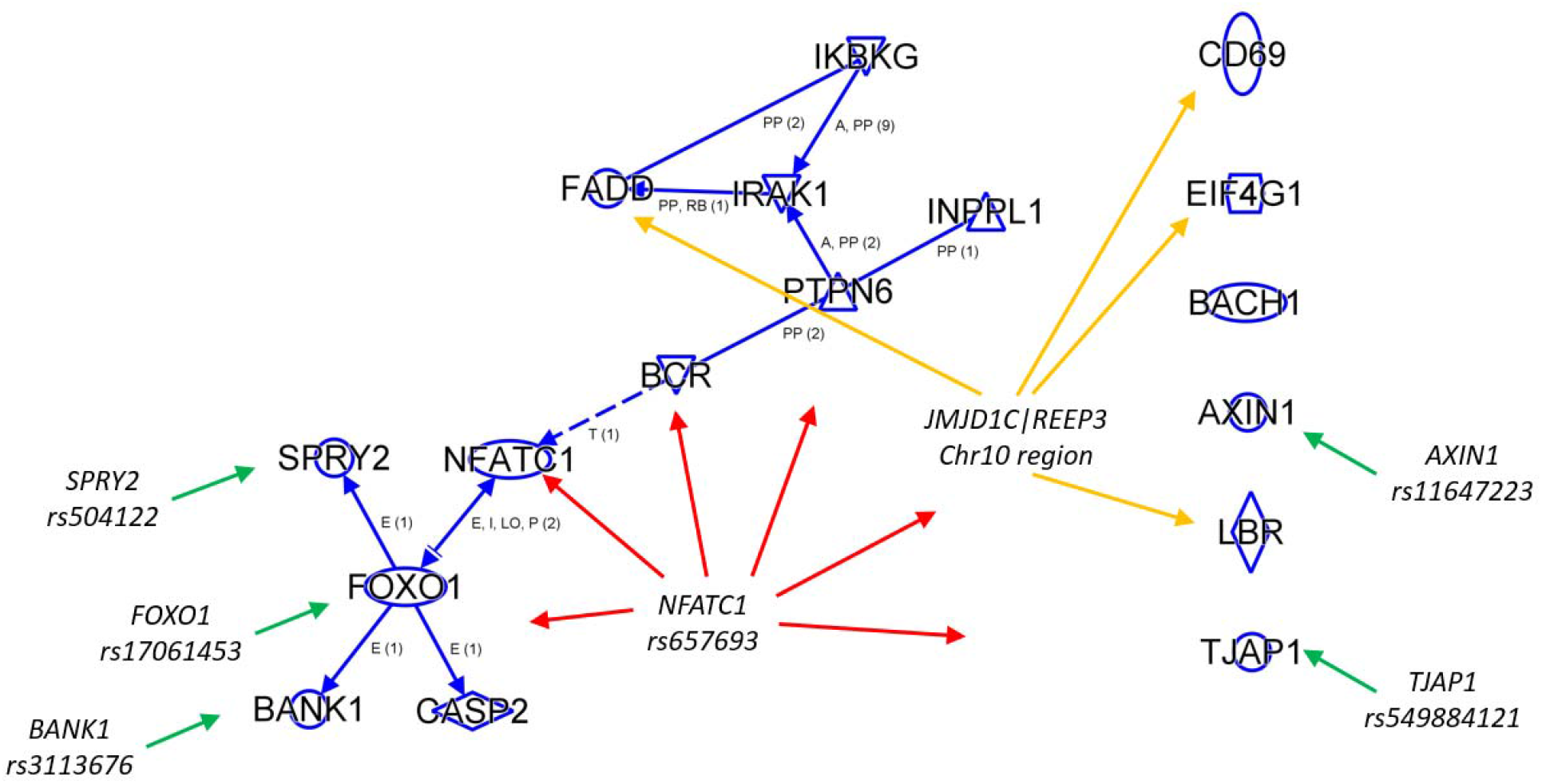
NFATC1 network. Protein-protein interactions obtained using Ingenuity Pathway Analysis (IPA)’s connect function with default settings (accessed 4 July 2023); Blue arrows indicate IPA interactions with the following abbreviations: A: activation, E: expression, I: inhibition, LO: localization, P: phosphorylation, PP: protein-protein interaction, RB: regulation of binding, T: transcription; Numbers in parentheses indicate multiple sources of evidence; Green arrows: rQTLs for the ratio of NFATC1 with the respective proteins at a *cis-*location; Orange: multiple variants on Chr10 around 65 MB associated with ratios between the listed proteins and NFATC1; Red: rs657693 at the *NFATC1* gene locus associated with all depicted proteins in a ratio with NFATC1 (details are in **Supplementary Table 4**).

To further explore the benefits of all-against-all ratios, we computed the associations of SNP rs12075 with all possible rations between 76 cytokines that are in the Olink panel. rs12075 (1:159175354:G:A) is an amino acid changing variant (c.125G>A, p.Gly42Asp) in the *atypical chemokine receptor 1* gene (*ACKR1* aka *DARC*). The glycine variant defines the Fy^a^ allele and the aspartate variant the Fy^b^ allele of the Duffy blood group system ^31^. DARC is clinically important as it is the entry point for the human malaria parasite *Plasmodium vivax.* Individuals with two copies of the FY^a^ allele or a silenced FY^b^ allele are resistant to *Plasmodium vivax* infection. A structural basis for DARC binding to *Plasmodium vivax’ Duffy-binding protein* involving the region around the p.Gly42Asp variant has been proposed ^32^.

Eleven cytokines associated in the UKB PPP GWAS at the ACKR1 locus (CCL2, CCL7, CCL8, CCL11, CCL13, CCL14, CCL17, CCL26, CXCL1, CXCL6, CXCL8; **Supplementary Table 3**). CCL7 and CCL8 protein levels increased with copy number of the Fy^a^ allele, while levels of the other nine cytokines decreased with that variant. DARC controls chemokine levels through promiscuous binding ^33^. The associations with these cytokines are thus matching the function of DARC. In our discovery study, rs12075 associated with three ratios (CCL13 / CCL8, CCL2 / CCL7, and CCL11 / CCL7; **Supplementary Table 4**), the strongest association of rs12075 was with the ratio between CCL8 and CCL13. Testing the association of rs12075 with all possible ratios between the 76 cytokines on the Olink panel implicated twelve additional cytokines (CCL3, CCL4, CCL14, CXCL11, CXCL12, HGF, IL7, PDGFA, TGFB1, THPO, TNFSF13, TNFSF14) in significant (p-gain > 10^10^) rQTLs (**Supplementary Table 8** and **Figure 7**). It goes beyond the scope of the present study to interpret these associations in further detail. The take-away message here is that using ratios we not only identified additional cytokines that associate with the Duffy blood type, but also suggest interactions between specific pairs of proteins, like CXCL6 that occurs in a significant ratio with CCL8, but not with CCL7. Especially at pleiotropic loci, where multiple proteins associate with a clinically relevant variant, it may be worthwhile to conduct this kind of all-against-all ratio analysis, using a subset of functionally related protein, as done in this example, or even extending the ratio analysis to the full protein panel.

**Figure 7:**
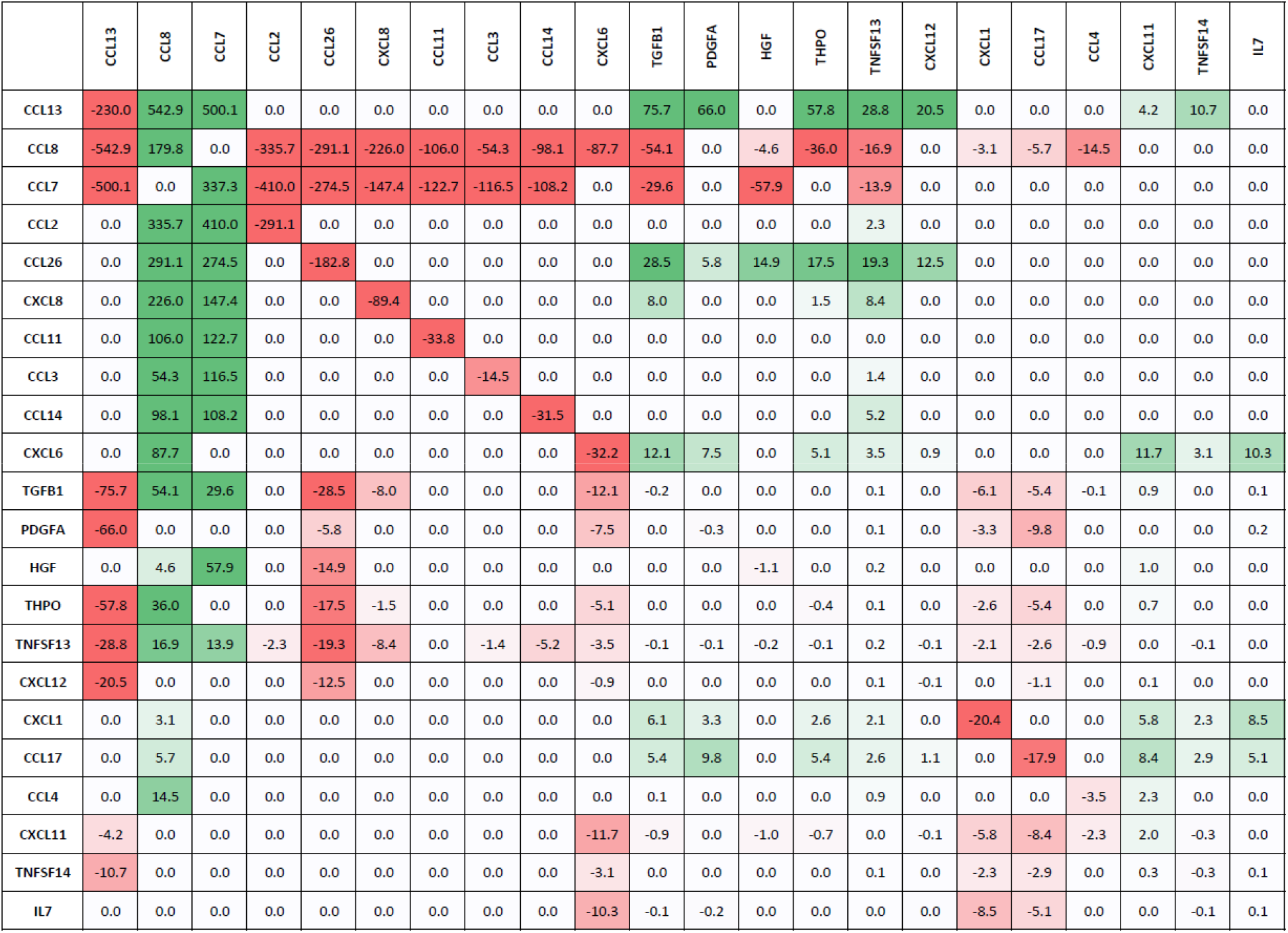
P-gain matrix for the association of rs12075 with all ratios between cytokines. SNP rs12075 (1:159175354:G:A) is an amino acid changing variant in ACKR1 aka DARC (c.125G>A, p.Gly42Asp) and defines the co-dominant Duffy blood type alleles Fy^a^ (Gly) and Fy^b^ (Asp). Limited to associations with -log10(p-value) > 10 or log10(p-gain) > 10 (full matrix in Supplementary Table 6); Values on the diagonal are -log10(p-value) for the single protein associations; Values in the off-diagonal cells are -log10(p-gain); The directionality of the associations with the copy number of the Fy^a^ allele are indicated by the sign and colored red (negative association) and green (positive association). Note that the -log10(p-value) for the ratios can be obtained by adding the log10(p-gain) of the ratio to the larger of the two -log10(p-value) of the single protein associations (full data in **Supplementary Table 8**).

## DISCUSSION

This is, to the best of our knowledge, the first GWAS at scale with ratios between blood circulating protein levels, using the recently released Olink proteomics data for almost 1,500 proteins measured in blood plasma of 54,000 participants of the UK Biobank. Using ratios, we observed increases in the strengths of association by up to several hundred orders of magnitude, involving two thirds of the proteins targeted by the Olink platform, increasing the strength of association at 16% of the pQTLs from the UKB PPP GWAS. We further reported novel cis-pQTLs for 13% of the 300 Olink proteins for which such a target-confirmatory QTL had not been identified so far and uncovered over 2,500 novel QTLs with ratios at loci that had not been highlighted by the UKB PPP GWAS using single protein levels, which corresponds to a 25% increase in the discovery rate.

We argue that ratios can account for unidentified genetic and/or non-genetic variance that is shared between the associated protein pairs (**Figure 5**). rQTL protein pairs were 7.2-fold enriched in known protein-protein interactions, demonstrating that they add substantial new information to hypothesis generation and providing a broad set of protein-protein relationships that can be mined using network pharmacology (NFATC1 example) ^34^ and systems immunology (ACKR1 example) ^35, 36^ approaches. We further reported selected examples that illustrate novel insights gained from using ratios, adding new information to established loci (GPX1 with IBD example) and identifying entirely novel loci (STK11 with AD example).

As we could only discuss a fraction of the biologically relevant findings in this paper, we freely share the full summary statistics of the GWAS using array-genotyped data and of the refinement using imputed genotype data, together with the corresponding Manhattan and regional association plots. These data represent a rich resource for biomedical hypothesis generation that complements the data generated by the UKB PPP GWAS and should be of particular value for pharmaceutical drug target development.

The following caveats apply: We analyzed only a subset of all possible ratios. Although the likelihood of finding a significant ratio association for a protein pair increased with the strength of their partial correlation, even for uncorrelated proteins this likelihood remains high (5%), supporting the testing of all possible ratio in future GWAS, if resources permit. Indeed, we hope that the present study shall motivate further methods development that could render all-against-all ratio testing computationally feasible, and maybe also more research into formal statistical methods that may generalize the analysis of combinations of quantitative traits as dependent variables in GWAS.

It should also be noted that affinity proteomics technologies have numerous limitations, such as effects of epitope changing variants, non-specific binding, and uncertainty about target specificity. However, the many biologically relevant associations that have been derived using data from the Olink and other affinity proteomics platforms suggest that these concerns are of minor relevance. The fact that we identified 39 novel *cis-*pQTLs provides further confirmatory evidence for the target specificity of their respective affinity binders.

Taken together, we hope to have demonstrated the benefits of analyzing ratios between protein levels at scale, an approach that we believe has already shown its benefits in the metabolomics field. Future work is needed to further speed up ratio associations, especially in the light of broadening proteomics panels and their increased application in large-scale cohorts. Also, further theoretical development and generalization of the concept of using ratios in more thorough statistical terms may be beneficial, as there are similarities in the approach to what could be conceived as “local Mendelian randomization”.

## METHODS

### Data sources

All data was obtained through the UKB RAP system on the DNAnexus platform (data dispensed on April 23, 2023; application id 43418) for samples satisfying the criterion “Number of proteins measured | Instance 0” is greater than “0” (https://biobank.ndph.ox.ac.uk/showcase/field.cgi?id=30900). Imputed genotypes ^37^ were extracted from BGEN files (https://biobank.ndph.ox.ac.uk/showcase/label.cgi?id=100319) using bgenix ^38^ and reformatted to text format (.raw) using plink ^39^ for further analysis in R. Phenotype data for age, sex, BMI, the first three genotype principal components, and the classification of genetic ethnic grouping were extracted using the DNAnexus cohort browser and the table downloader app (https://ukbiobank.dnanexus.com/landing). Details on genomics and proteomics data QC and preprocessing are available in the accompanying UK Biobank resource files (available at the respective showcase links given above).

The downloaded proteomics data set comprised NPX values for 1,463 proteins for 52,749 participants. NPX values correspond to relative protein concentrations and are reported on a log-scale. Data analysis was restricted to 52,705 samples that were collected at baseline (instance 0). Samples were split into a discovery set of 43,509 samples identified as Caucasian based on the genetic ethnic grouping variable and a replication set of 9,196 ethnically diverse samples (https://biobank.ndph.ox.ac.uk/showcase/field.cgi?id=22006). A total of 5,717 unique variants corresponding to 10,248 pQTLs of the UKB PPP GWAS were analyzed. These pQTLs were obtained from Supplementary Table 6 of Sun *et al*. ^6^.

### Graphical Gaussian model

Partial correlations were computed using the R function ggm.estimate.pcor from the package GeneNet ^40^. All baseline samples were used for this step. As this analysis does not allow for the presence of missing values, samples with more than 20% missing protein values were removed (N=1,840), followed by proteins that were missing in more than 20% of the samples (N=3). The remaining missing data points were imputed to minimum (N=37,419). A total of 11,936 partial correlations were identified at a Bonferroni significance cut-off p-value of 4.7×10^-8^. The smallest r^2^ at this level was 0.00176.

### Statistical analysis

Linear models with inverse-normal scaled proteomics data (NPX values) as dependent variables and genotype, age, sex, and the first three genotype principal components were computed using the R function “lm”. For ratios, inverse-normal scaled differences between the two NPX values were used, based on the relation log(A/B) = log(A) - log(B) and the NPX values representing protein levels on a log-scale. The p-gain for associations with ratios between two protein traits was computed as the smaller of the two p-values for the individual trait associations divided by the p-value for the ratio association ^41^. Log10-scaled p-values and p-gains were used throughout to avoid numeric overflows and rounding of small p-values to zero.

For all pairs of proteins with a Bonferroni significant partial correlation all variants that were associated with at least one of the two proteins in a pQTL in the UKB PPP GWAS were identified. For these variant – protein pairs the single protein and ratio association statistics were computed using the discovery and replication samples separately.

### GWAS analysis

The GWAS on 2,821 ratios was conducted using plink2 ^39^ on the UKB RAP platform hosted by DNAnexus with the --glm option, using age, sex, and the first the genoPCs as covariates. We used the array-genotyped UKB data with the following variant filtering options: --geno 0.1, --hwe 1e-15, --mac 100, --maf 0.01, --mind 0.1.

### Other sources of data

The STRING database of proteins and their functional interactions was used to identify known relationships between proteins ^42^. The database was downloaded from https://string-db.org/cgi/download (version 11.5, accessed 5 May 2023). Annotated cytokine and cytokine-receptor pairs were downloaded from CytokineLink ^26^ (https://github.com/korcsmarosgroup/CytokineLink, accessed 30 June 2023). Drug target development status was obtained from NCBI Pharos ^15^ (accessed 11 July 2023). Variants were annotated with PhenoScanner API ^16^ using proxies based on EUR LD r^2^>0.8 (accessed 5 May - 11 July 2023). LocusZoom ^43^ was used to generate regional association plots with LD annotation (EUR population). GeneCards ^44^ was used to obtain general information about the associated genes and proteins.

## AUTHOR STATEMENTS

### Data availability statement

All analyzed UK Biobank data was obtained through the UKB RAP system under application reference number 43418 and is accessible upon application online via https://biobank.ndph.ox.ac.uk/showcase/. Full summary statistics for all 2,821 GWAS with ratios using the array-genotyped UKB data shall be made available via the GWAS catalogue. Summary statistics using imputed UKB genotype data for the regions (+/-500kb) around the 8,462 rQTLs discovered in the GWAS are available on FigShare (doi:10.6084/m9.figshare.23695398), both for the discovery and the replication cohort. Manhattan and regional association plots based on this data are available in PDF format on FigShare at the same URL.

### Ethics statement

All data and samples were collected by UK Biobank following all relevant ethical guidelines and procedures and were shared with the UKB users under rules reviewed by the UK Biobank ethics committee and board.

### Code availability statement

Only publicly available software was used in the data analysis (R and Rstudio).

## Acknowledgements

We thank all UK Biobank participants for their contribution.

## Author contributions

K.S. conceived the study, conducted the data analyses and wrote the paper.

## SUPPLEMENTARY TABLES

Supplementary tables are provided in EXCEL format.

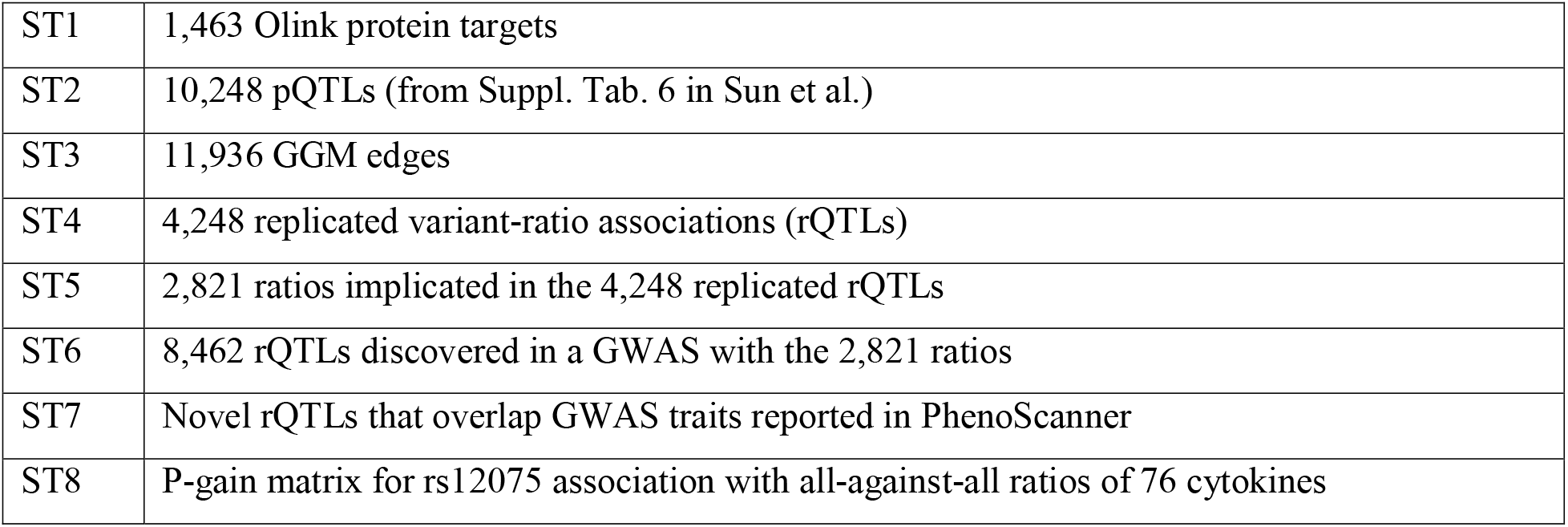

## SUPPLEMENTARY FIGURES

**Supplementary Figure 1:**
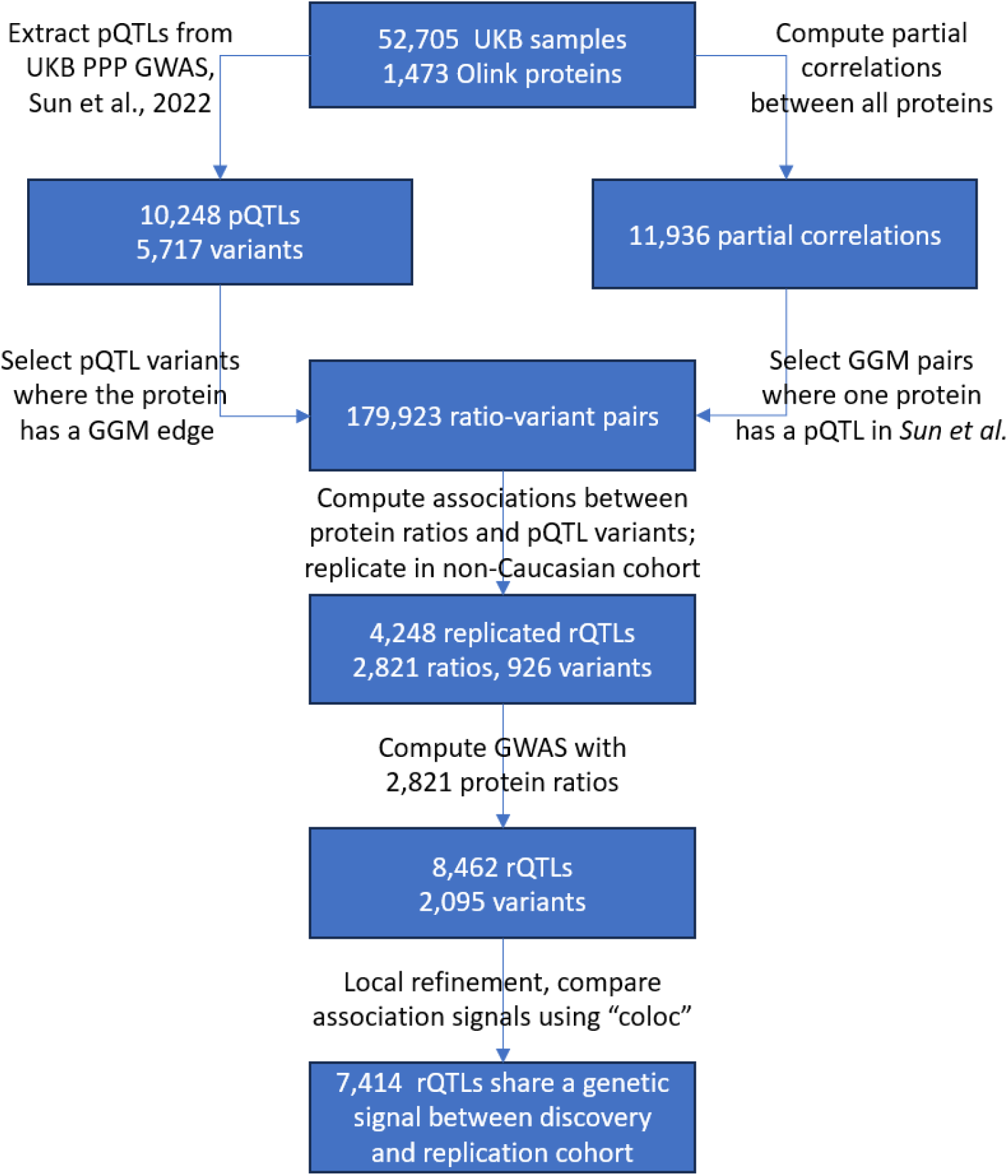
Flowchart of the study.

**Supplementary Figure 2:**
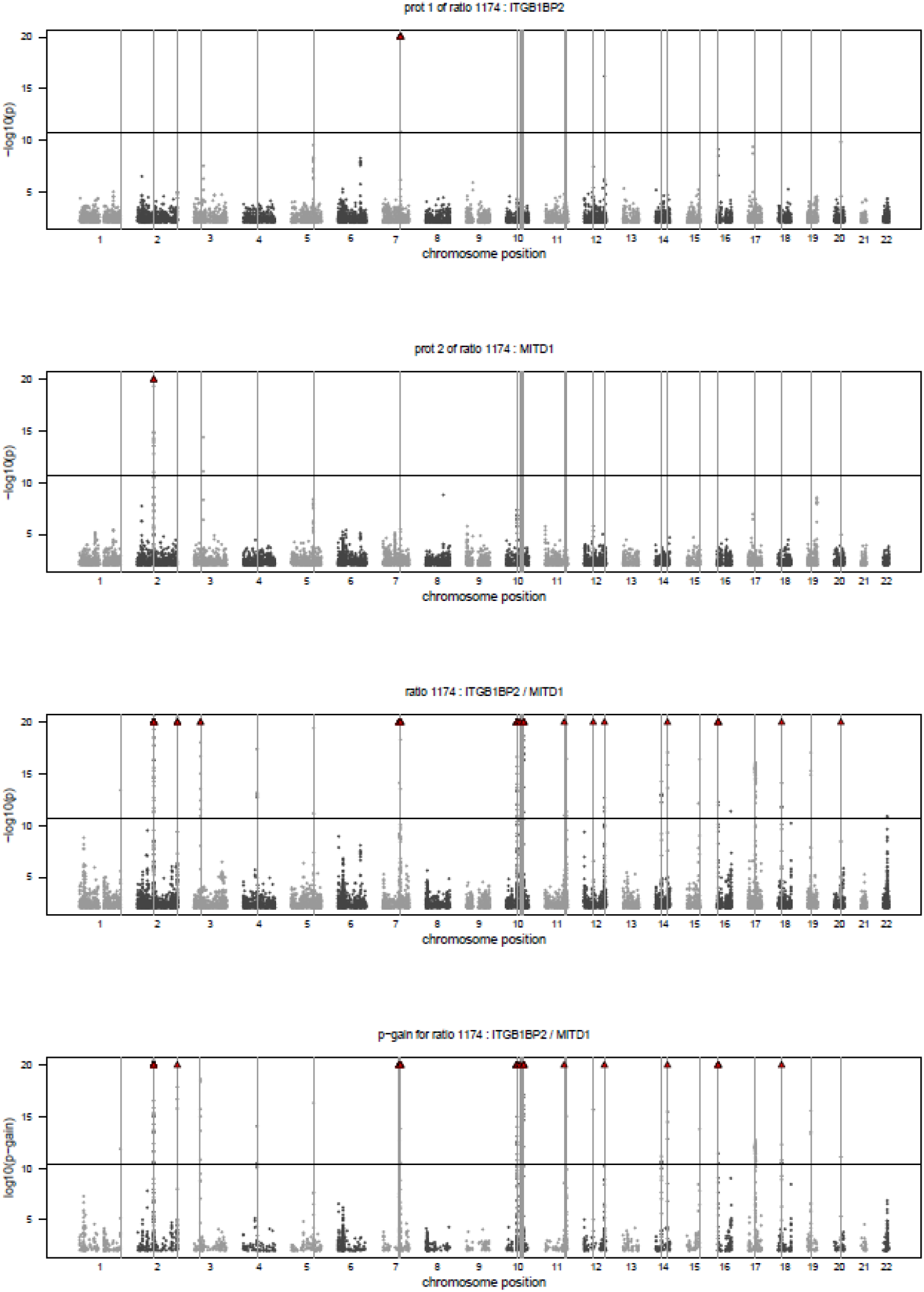
Example of Manhattan plots for a ratio. This Figure explains how by using ratios novel genetic loci can be uncovered. Plotted are the associations of the two individual proteins (ITGB1BP2 and MTD1), the ratio (ITGB1BP2 / MTD1), and the p-gain of the ratio using array genotype data; Similar Manhattan plots are available in PDF format for the 2,821 ratios on FigShare (doi:10.6084/m9.figshare.23695398); The full GWAS summary statistics will be deposited with the GWAS catalogue.

**Supplementary Figure 3:**
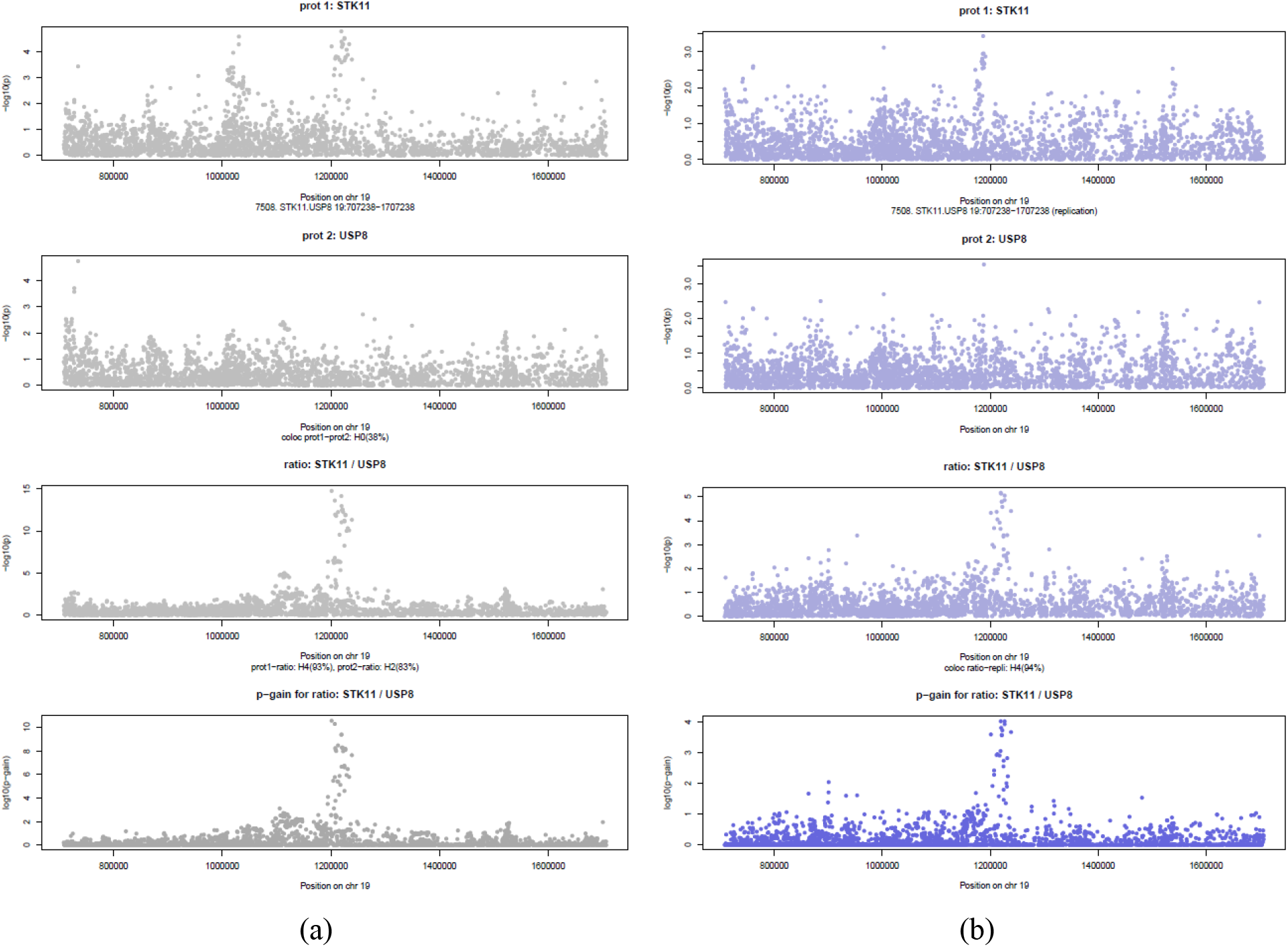
Example of regional association plots for rQTLs. This Figure explains how by using ratios a genetic signal can emerge from the noise. Plotted are the associations of the two individual proteins (STK11 and USP8), the ratio (STK11 / USP8), and the p-gain of the ratio using imputed genotype data +/-500kb around the variant rs3764640 for the discovery (a) and the replication cohort (b); The subtitles indicate the most likely *coloc* hypotheses regarding the similarity between the relevant genetic signals; Similar regional association plots together with the full summary statistics used in these plots for 8,462 rQTLs are available in PDF format on FigShare (doi:10.6084/m9.figshare.23695398).

## SUPPLEMENTARY DATA

The following items are available on FigShare (doi:10.6084/m9.figshare.23695398):

| Item | Description |
| --- | --- |
| Local association data | Full summary statistics for +/- 500kb regional refinements around 8,462 rQTL lead SNPs using imputed genotype data. |
| Manhattan plots | Manhattan plots for the GWAS (PDF format) |
| Regional association plots | Regional association plots for local refinements (PDF format) |

## REFERENCES

1. Suhre, K., McCarthy, M.I. & Schwenk, J.M. Genetics meets proteomics: perspectives for large population-based studies. Nat Rev Genet 22, 19–37 (2021).

2. Sun, B.B. et al. Genomic atlas of the human plasma proteome. Nature 558, 73–79 (2018).

3. Folkersen, L. et al. Genomic and drug target evaluation of 90 cardiovascular proteins in 30,931 individuals. Nat Metab 2, 1135–1148 (2020).

4. Emilsson, V. et al. Co-regulatory networks of human serum proteins link genetics to disease. Science 361, 769–773 (2018).

5. Suhre, K. et al. Connecting genetic risk to disease end points through the human blood plasma proteome. Nat Commun 8, 14357 (2017).

6. Sun, B.B. et al. Genetic regulation of the human plasma proteome in 54,306 UK Biobank participants. bioRxiv, 2022.06.17.496443 (2022).

7. Suhre, K. & Gieger, C. Genetic variation in metabolic phenotypes: study designs and applications. Nat Rev Genet 13, 759–69 (2012).

8. Kastenmuller, G., Raffler, J., Gieger, C. & Suhre, K. Genetics of human metabolism: an update. Hum Mol Genet 24, R93–r101 (2015).

9. Krumsiek, J. et al. Mining the unknown: a systems approach to metabolite identification combining genetic and metabolic information. PLoS Genet 8, e1003005 (2012).

10. Krumsiek, J., Suhre, K., Illig, T., Adamski, J. & Theis, F.J. Gaussian graphical modeling reconstructs pathway reactions from high-throughput metabolomics data. BMC Syst Biol 5, 21 (2011).

11. Petersen, A.K. et al. On the hypothesis-free testing of metabolite ratios in genome-wide and metabolome-wide association studies. BMC Bioinformatics 13, 120 (2012).

12. Suhre, K. et al. Human metabolic individuality in biomedical and pharmaceutical research. Nature 477, 54–60 (2011).

13. Shin, S.Y. et al. An atlas of genetic influences on human blood metabolites. Nat Genet 46, 543–550 (2014).

14. Surendran, P. et al. Rare and common genetic determinants of metabolic individuality and their effects on human health. Nat Med 28, 2321–2332 (2022).

15. Sheils, T.K., et al. TCRD and Pharos 2021: mining the human proteome for disease biology. Nucleic Acids Research 49, D1334–D1346 (2020).

16. Staley, J.R. et al. PhenoScanner: a database of human genotype-phenotype associations. Bioinformatics 32, 3207–3209 (2016).

17. Hollingworth, P. et al. Genome-wide association study of Alzheimer’s disease with psychotic symptoms. Molecular Psychiatry 17, 1316–1327 (2012).

18. Lu, Z. & Hunter, T. Degradation of activated protein kinases by ubiquitination. Annu Rev Biochem 78, 435–75 (2009).

19. Liu, J.Z. et al. Association analyses identify 38 susceptibility loci for inflammatory bowel disease and highlight shared genetic risk across populations. Nat Genet 47, 979–986 (2015).

20. Häuser, F. et al. Inflammatory bowel disease (IBD) locus 12: is glutathione peroxidase-1 (GPX1) the relevant gene? Genes & Immunity 16, 571–575 (2015).

21. Wallace, C. A more accurate method for colocalisation analysis allowing for multiple causal variants. PLOS Genetics 17, e1009440 (2021).

22. Giambartolomei, C. et al. Bayesian test for colocalisation between pairs of genetic association studies using summary statistics. PLoS Genet 10, e1004383 (2014).

23. Illig, T. et al. A genome-wide perspective of genetic variation in human metabolism. Nat Genet 42, 137–41 (2010).

24. Lagace, T.A. et al. Secreted PCSK9 decreases the number of LDL receptors in hepatocytes and in livers of parabiotic mice. J Clin Invest 116, 2995–3005 (2006).

25. Lagace, T.A. PCSK9 and LDLR degradation: regulatory mechanisms in circulation and in cells. Curr Opin Lipidol 25, 387–93 (2014).

26. Olbei, M. et al. CytokineLink: A Cytokine Communication Map to Analyse Immune Responses-Case Studies in Inflammatory Bowel Disease and COVID-19. Cells 10(2021).

27. Macian, F. NFAT proteins: key regulators of T-cell development and function. Nat Rev Immunol 5, 472–84 (2005).

28. Lee, J.U., Kim, L.K. & Choi, J.M. Revisiting the Concept of Targeting NFAT to Control T Cell Immunity and Autoimmune Diseases. Front Immunol 9, 2747 (2018).

29. Gao, R., Zhang, Y., Zeng, C. & Li, Y. The role of NFAT in the pathogenesis and targeted therapy of hematological malignancies. European Journal of Pharmacology 921, 174889 (2022).

30. Metzelder, S.K. et al. NFATc1 as a therapeutic target in FLT3-ITD-positive AML. Leukemia 29, 1470–1477 (2015).

31. King, C.L. et al. Fy(a)/Fy(b) antigen polymorphism in human erythrocyte Duffy antigen affects susceptibility to Plasmodium vivax malaria. Proc Natl Acad Sci U S A 108, 20113–8 (2011).

32. Moskovitz, R. et al. Structural basis for DARC binding in reticulocyte invasion by Plasmodium vivax. Nat Commun 14, 3637 (2023).

33. Nibbs, R.J.B. & Graham, G.J. Immune regulation by atypical chemokine receptors. Nature Reviews Immunology 13, 815–829 (2013).

34. Hopkins, A.L. Network pharmacology: the next paradigm in drug discovery. Nature Chemical Biology 4, 682–690 (2008).

35. Altan-Bonnet, G. & Mukherjee, R. Cytokine-mediated communication: a quantitative appraisal of immune complexity. Nature Reviews Immunology 19, 205–217 (2019).

36. Dhillon, B.K., Smith, M., Baghela, A., Lee, A.H.Y. & Hancock, R.E.W. Systems Biology Approaches to Understanding the Human Immune System. Front Immunol 11, 1683 (2020).

37. Bycroft, C. et al. The UK Biobank resource with deep phenotyping and genomic data. Nature 562, 203–209 (2018).

38. Band, G. & Marchini, J. BGEN: a binary file format for imputed genotype and haplotype data. bioRxiv, 308296 (2018).

39. Purcell, S. et al. PLINK: a tool set for whole-genome association and population-based linkage analyses. Am J Hum Genet 81, 559–75 (2007).

40. Schäfer, J. & Strimmer, K. A shrinkage approach to large-scale covariance matrix estimation and implications for functional genomics. Stat Appl Genet Mol Biol 4, Article32 (2005).

41. Petersen, A.K. et al. On the hypothesis-free testing of metabolite ratios in genome-wide and metabolome-wide association studies. BMC Bioinformatics 13, 120 (2012).

42. Szklarczyk, D., et al. STRING v11: protein-protein association networks with increased coverage, supporting functional discovery in genome-wide experimental datasets. Nucleic Acids Res 47, D607–d613 (2019).

43. Boughton, A.P. et al. LocusZoom.js: interactive and embeddable visualization of genetic association study results. Bioinformatics 37, 3017–3018 (2021).

44. Stelzer, G. et al. The GeneCards Suite: From Gene Data Mining to Disease Genome Sequence Analyses. Curr Protoc Bioinformatics 54, 1.30.1–1.30.33 (2016).

